# Ubiquitin protease Ubp1 cooperates with Ubp10 and Ubp12 to revert Lysine-164 PCNA ubiquitylation at replication forks

**DOI:** 10.1101/2023.10.31.564928

**Authors:** Javier Zamarreño, Avelino Bueno, María P. Sacristán

## Abstract

Proliferating cell nuclear antigen (PCNA) is essential for the faithful duplication of eukaryotic genomes. PCNA orchestrates events necessary to deal with threats to genomic integrity, such as the DNA damage tolerance (DDT) response. DDT is a mechanism by which eukaryotic cells bypass replication-blocking lesions to prevent replisome instability. DDT pathways are regulated by the ubiquitylation of PCNA and the consequent recruitment of specialized polymerases and mechanisms able to guarantee the continuity of replication. We have previously described that the deubiquitylases Ubp10 and Ubp12 associate with replication forks and modulate DDT events by reverting the ubiquitylation of PCNA in *Saccharomyces cerevisiae*. The results of this study unveil Ubp1 as a new PCNA deubiquitylase, which cooperates with Ubp10 and Ubp12 in the regulation of DDT during DNA replication. Ubp1 is known as a cytoplasmic protein, however, we found that it also localizes to the nucleus where it binds to chromatin and associates with DNA replication forks. In addition, Ubp1 interacts with and deubiquitylates PCNA. The ablation of Ubp1, Ubp10, and Ubp12 enhances both the accumulation of ubiquitylated PCNA and the DNA replication defects observed in cells depleted for Ubp10 and Ubp12, supporting a cooperative role among the three enzymes.

**IMPORTANCE:** PCNA ubiquitylation regulates DDT mechanisms to bypass genotoxic lesions during replication and that PCNA deubiquitylation is required to limit the extent of bypass events. This study shows that *Saccharomyces cerevisiae* PCNA is ubiquitylated during an unperturbed S-phase progression and that three ubiquitin proteases (Ubp1, Ubp10, and Ubp12) work together facilitating DNA replication by efficiently controlling ubiquitylation of PCNA at replication forks.

## INTRODUCTION

DNA replication is one of the most fundamental biological functions in life, as it faithfully copies genomes and propagates genetic material from generation to generation. To this end, replisomes, complex molecular machines assembled at specific replication initiation sites, initiate and carry out DNA synthesis under rigorous and overlapping regulatory mechanisms that overcome potential replication obstacles to ensure genetic stability and cell viability. The replication obstacles comprise both endogenous and exogenous DNA damage to which cells are exposed. During the S-phase, cells are particularly sensitive to DNA lesions due to the highly accurate nature of replicative DNA polymerases, thus failing to accommodate damaged nucleotides as templates and, consequently, blocking replication forks. Persistent replication stalling results in detrimental effects on genomic stability and cell viability.

To cope with DNA lesions during replication, organisms evolved DNA damage tolerance (DDT) mechanisms that allow them to bypass the damage ensuring a timely coordination between replication fork progression and DNA repair (1–3). DDT is exerted through two major pathways: error-prone translesion synthesis (TLS) and error-free template switching (TS), which involve different mechanisms (3). TLS DNA polymerases are specialized, evolutionary conserved, alternative DNA polymerases capable of replicating across damaged bases and bypass lesions, although they are error-prone and potentially mutagenic (4). In contrast, TS-based DDT mechanisms involve the pairing of a blocked nascent strand with its sister chromatid to copy an intact base, providing an error-free pathway (reviewed by (5)).

Proliferating cell nuclear antigen (PCNA) plays a key role in DNA synthesis and DDT (6–8). PCNA constitutes a moving platform that accurately recruits different factors essential for replication or DDT. Three monomers of PCNA form a ring-shaped homo-trimer that encircles DNA and recruits replicative DNA polymerases to carry out high-fidelity DNA synthesis (9). Since PCNA lacks enzymatic activity, it exerts its functions through numerous protein-protein interactions, which are regulated by post-translational modifications (7, 10, 11). An essential mechanism in the regulation of DDT is ubiquitylation (7). When the replication machinery confronts DNA lesions, replication forks stall, and PCNA is mono-ubiquitylated at lysine residue (K) 164 by the evolutionary conserved RAD6/RAD18 (E2/E3) ubiquitin ligase complex (7, 12, 13). This modification changes the PCNA association from high-fidelity to low-fidelity polymerases (14–16), promoting the error-prone TLS DDT pathway (17–21), which is essential to prevent replication gaps that constitute a high risk of tumorigenesis (22). Furthermore, the addition of Lys^63^-linked ubiquitin residues to mono-ubPCNA^K164^ by the Rad5/Mms2/Ubc13 PCNA-ubiquitin ligase complex (2, 23) results in polyubiquitinated PCNA, which promotes the error-free TS DDT pathway and also prevents genomic instability and tumorigenesis (24, 25).

Low-fidelity TLS polymerases prevents persistent replication stalling, but their activity also increases the risk of introducing mutations opposite to DNA lesions, potentially leading to tumorigenesis (7, 13). Moreover, lesion bypass through TS mechanisms is error-free, but it also implies certain risks for cells due to the formation of structures between sister chromatids that hinder chromosome segregation (5). Hence, both TLS and TS DTT pathways should be limited to minimize their deleterious cellular side effects. In fact, PCNA-deubiquitylation processes have been involved in the control of DDT during DNA synthesis to support normal replication rates and prevent mutagenesis (26, 27).

It has been proposed that specialized PCNA-ubiquitin proteases, capable of removing ubiquitin residues conjugated to PCNA-K^164^, suppress DDT events and prevent genomic instability. The mammalian deubiquitinating enzymes (DUBs) Usp1, Usp7, and Usp10 revert PCNA ubiquitylation caused in response to DNA damage (26, 28–31). In human cells, the loss of USP1 induces aberrant PCNA monoubiquitylation leading to enhanced recruitment of error-prone TLS polymerases, which destabilize replication forks in cells lacking the homologous recombination factor BRCA1 (31). Moreover, the knockdown of USP1 in 293T human cells increases mutagenesis levels (28, 31). The ubiquitin proteases Ubp2, Ubp12, Ubp15, and Ubp16 cooperate to revert PCNA^K164^ ubiquitylation in the fission yeast *Schizosaccharomyces pombe* (32). In the case of the budding yeast *Saccharomyces cerevisiae*, the ubiquitin protease Ubp10 was initially identified as a PCNA-DUB that removes DNA damage- and replicative stress-dependent PCNA ubiquitylation (33). More recently, Ubp10 together with Ubp12 were found to be key enzymes that limit the extent of DDT processes during the progression of exogenously unperturbed S phase (27).

In this study, we analyzed the role of Ubp1, which is one of the 17 ubiquitin-specific proteases of the USP family in *S. cerevisiae* (34). *UBP1* gene codifies for two Ubp1 forms, a longer membrane-anchored form and a shorter soluble one (35). Ubp1, in particular the membrane-anchored form, has a well-studied role in the regulation of the endoplasmic reticulum-associated protein degradation pathway as a ubiquitin-specific protease of the Hrd1 protein (36). Moreover, the shorter form of Ubp1 has been involved in endocytosis (35). Here we present results supporting that Ubp1 is a ubiquitin-specific protease that also removes ubiquitin moieties from ubiquitylated PCNA in budding yeast. We observed that a fraction of Ubp1 localizes into the nucleus, interacts with PCNA, and associates with replication forks. Deletion of Ubp1 in combination with depletion of PCNA-DUBs, Ubp10 and Ubp12, increases and stabilizes the levels of ubiquitylated PCNA throughout an unperturbed S-phase. Moreover, *ubp1Δ ubp10Δ ubp12Δ* triple mutant cells exhibit a strong S phase progression defect, higher than the one observed in Ubp10/Ubp12-ablated cells, which is rescued by retention of Ubp1 in the nucleus. Finally, by 2D gels analysis, we found that Ubp1 contributes to solving transient TS-dependent replication structures generated upon replication stress. Altogether, our data suggest that Ubp1 contributes to proper S phase progression by cooperating in deubiquitylation-mediated regulation of PCNA. This study brings a new piece of knowledge about the still enigmatic processes of the regulation of PCNA in *S. cerevisiae* and contributes to a better understanding of the complex regulation of DNA replication to preserve genomic integrity and cell viability.

## RESULTS

### Ubp1 cooperates with Ubp10 and Ubp12 in PCNA deubiquitylation to regulate S phase progression

We have previously reported that in the budding yeast *S. cerevisiae,* the ubiquitin proteases Ubp10 and Ubp12 deubiquitylate PCNA during the S phase to regulate the DDT response allowing a processive DNA replication in unperturbed cycling cells. The lack of Ubp10 alone already causes a significant slow S phase progression, although it is the lack of both Ubp10 and Ubp12 proteases that is necessary to detect by immunoblotting the accumulation of ubiquitylated PCNA forms in asynchronous cell cultures (27).

To understand the participation of these two ubiquitin proteases in the dynamic ubiquitylation/deubiquitylation-dependent regulation of PCNA during DNA replication, the ubiquitylation state of PCNA in *ubp10Δ ubp12Δ* double mutant cells during S phase progression was analyzed by immunoblot. Cells were synchronized in G1 by treatment with α-factor and then released into fresh medium to allow progression through the S phase (Figure 1A). Samples were taken every 10 minutes for 2 hours and genome replication was followed by fluorescence-activated cell sorting (FACS). As expected, no ubiquitylated forms of PCNA were detected in wild-type cells. However, PCNA appeared ubiquitylated between the 50- and 90-minutes time points after α-factor release in the *ubp10Δ ubp12Δ* double mutant (Figure 1B). The accumulation of ubiquitylated PCNA forms correlated with a significant delay in the S phase progression of these cells (Figure 1C), previously observed in cells lacking Ubp10 alone (27), suggesting an important role of PCNA deubiquitylation in supporting normal replication rates. Surprisingly, however, ablation of the two known PCNA-DUBs, Ubp10 and Ubp12, did not fully prevent deubiquitylation of PCNA, and the ubiquitylated forms disappear at later replication time points (Figure 1B), implying that additional PCNA-DUBs are involved in this process.

**Figure 1.**
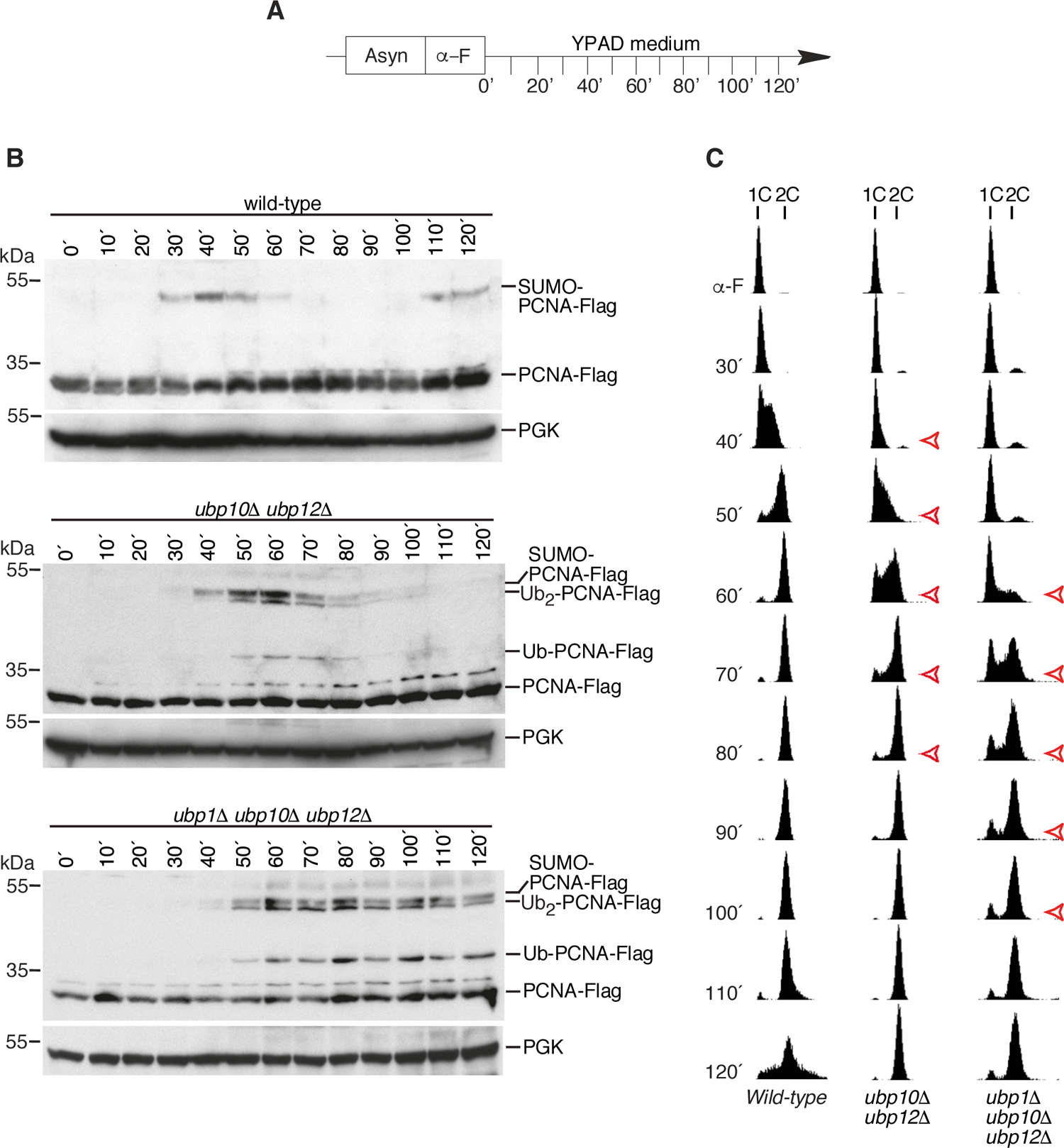
The lack of Ubp1 in combination with Ubp10 and Ubp12 causes a marked synergistic delay in S phase progression and a constant accumulation of ubiquitylated PCNA forms (A) Experimental design. Exponentially growing cultures of wild-type, *ubp1011. ubp1211* and *ubp111. ubp1011. ubp1211.* strains expressing PCNA-Flag fusion protein were synchronized at G1 by incubation with α-factor and then released into fresh yeast complex medium (YPAD). Samples were taken at indicated intervals and processed for FACS and immunoblot analysis. (B) Protein extracts were processed for immunoblotting with anti-Flag antibodies. 3-phosphoglycerate kinase (PGK) was used as a loading control. (C) DNA content analysis by FACS shows the progression of genome replication from α-factor synchronization to the 120 minutes time point after release. Red arrows indicate time points at which mutants show a replication delay. Two biological replicates were performed, and a representative experiment is shown.

Based on our previous biochemical screenings performed to identify the ubiquitin proteases involved in PCNA regulation (33), we analyzed cells lacking each of the different known DUBs in combination with the deletion of *UBP10* and *UBP12*, looking for increased levels of ubiquitylated PCNA in each triple mutant generated. We found that deletion of *UBP1,* another of the 17 known ubiquitin specific protease genes in *S. cerevisiae*, together with the absence of *UBP10* and *UBP12* resulted in a significant increase in ubiquitylated PCNA forms compared to that observed in *ubp10Δ ubp12Δ* double mutant cells asynchronously growing (Figure S1), strongly suggesting that Ubp1 was also involved in the deubiquitylation of PCNA.

We next analyzed the pattern of PCNA ubiquitylation during S phase progression in the triple mutant *ubp1Δ ubp10Δ ubp12Δ*. As shown in Figure 1B, the lack of *UBP1* in combination with the absence of *UBP10* and *UBP12* caused the stabilization of ubiquitylated PCNA forms from 50 minutes after α-factor release until the end of the time course analysis. Interestingly, the lack of deubiquitylation of PCNA in the triple mutant correlated with a remarkably extended S phase, wider than the one observed in Ubp10/Ubp12-ablated cells (Figure 1C). These data point to Ubp1 as a novel PCNA deubiquitylase collaborating with Ubp10 and Ubp12 in the regulation of S phase progression.

### A nuclear soluble form of Ubp1 has a role in PCNA deubiquitylation during unperturbed DNA replication

Ubp1 is a ubiquitin protease involved in the regulation of endoplasmic reticulum-associated protein degradation and vesicle trafficking pathways (35, 36). Ubp1 has been described as a cytoplasmic protein with two different isoforms originated from two transcription initiation sites (methionine residues 01 and 67) (35) (Figure 2A). The two isoforms correspond to the full-length isoform, which is membrane-anchored (mUbp1) through an N-terminal transmembrane (TM) segment and localized to the endoplasmic reticulum, and a shorter one that lacks the TM domain and is soluble (sUbp1) (35). The above results, according to which the lack of Ubp1 can increase the PCNA ubiquitylation state, led us to hypothesize the existence of a not-yet-described nuclear population of Ubp1 that might collaborate in the regulation of PCNA ubiquitylation. To address this point, we focused on the localization of Ubp1, generating two constructs in which Ubp1 was labeled with the fluorescent GFP epitope linked or not to a nuclear exclusion signal (NES) that efficiently prevents its potential nuclear location (32) (Figure 2A). As expected, the Ubp1-GFP fusion protein was distributed all over the cell, a pattern unable to confirm a clear nuclear location. However, Ubp1-GFP:NES showed a specific cytoplasmic location, clearly excluded from the DAPI signal (Figure 2B) suggesting that Ubp1 is also a nuclear protein. Next, we checked the accumulation of PCNA ubiquitylation in Ubp1-GFP:NES mutants in combination with the ablation of Ubp10 and Ubp12 under asynchronous growing conditions. As shown in Figure 2C, the exclusion of Ubp1 from the nucleus recapitulates the *UBP1* ablation phenotype. PCNA is widely ubiquitylated in response to DNA damage or replication stress. Therefore, we tested the ability of *ubp1011. ubp1211. Ubp1-GFP:NES* triple mutant to accumulate ubiquitylated PCNA forms upon the induction of replicative stress by treatment with the alkylating agent methyl methanesulphonate (MMS) or the ribonucleotide reductase inhibitor hydroxyurea (HU). Using a rabbit polyclonal antibody that detects PCNA and its monoubiquitylated forms (Figure S2) we confirmed that *ubp1011. ubp1211.* double mutant cells accumulate higher levels of PCNA ubiquitylation than the wild-type cells (27), and Figure 2C. Moreover, similar to what was observed for unperturbed cycling cells, the combination of *ubp111.* and *ubp1011. ubp1211.* determined a synergistic increase in ubiquitylated PCNA levels in both HU- and MMS-treated cells, which was also observed in the *ubp1011. ubp1211. ubp1-GFP:NES* mutant (Figure 2C; compare lanes 2 and 3-4). These data indicate the existence of a nuclear population of Ubp1 that plays a role in PCNA deubiquitylation.

**Figure 2.**
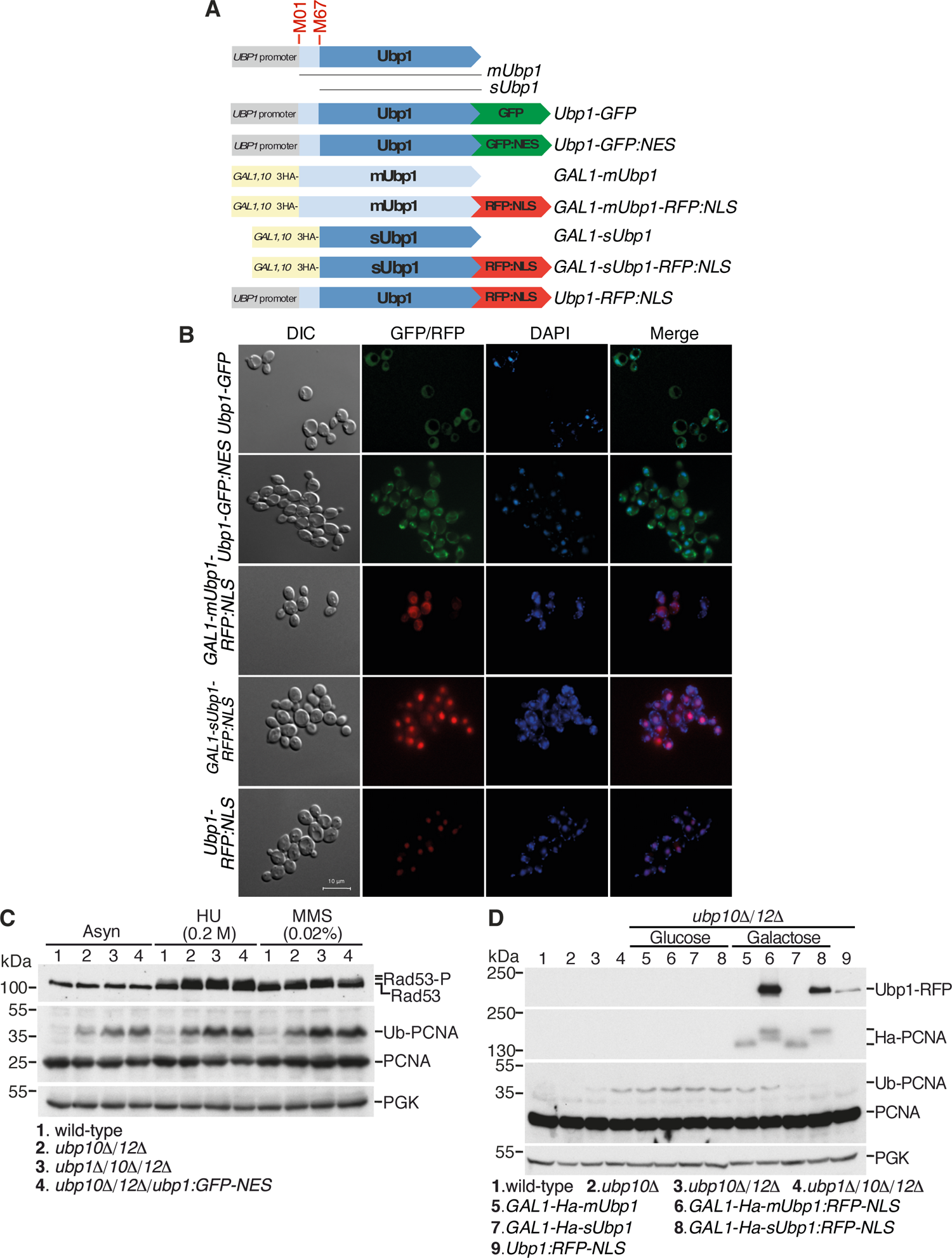
A soluble nuclear population of ubiquitin protease Ubp1 affects the ubiquitination state of PCNA (A) Scheme of the chimeric Ubp1 enzyme constructs used in B, C, and D. The *UBP1* gene shows two transcription initiation sites (M01 and M67), from which two different isoforms are originated (*mUbp1* and *sUbp1*). M, methionine residue; NES, Nuclear Exclusion Signal; NLS, Nuclear Localization Signal. (B) Fluorescence microscopy analyses of Ubp1 constructs labeled with GFP or RFP fluorescence proteins as indicated in A. Cells were stained with DAPI to visualize DNA. Bar, 10μm. (C) Analysis of ubiquitylated PCNA levels of the indicated strains under both untreated and treated conditions. Asynchronously growing cells (Asyn) were treated with 0.2 M HU or 0.02% MMS for 1 h. Total protein was extracted, resolved by 10% SDS-PAGE gels and immunoblotted with polyclonal anti-PCNA antibody. Rad53 phosphorylation was used to test checkpoint activation upon treatments. PGK immunoblot was used as a loading control. (D) Analysis of ubiquitylated PCNA levels of the indicated strains growing under unperturbed conditions. *ubp1011. ubp1211.* strain also containing *GAL1-mUbp1, GAL1-mUbp1:RFP-NLS, GAL1-sUbp1,* and *GAL1-sUbp1:RFP-NLS* constructs were incubated in glucose or galactose as unique carbon sources. Expression of the different chimeric Ubp1 forms was either repressed by adding glucose or induced by the addition of galactose to the medium. Note that when glucose was present, the *GAL1-mUbp1, GAL1-mUbp1:RFP-NLS, GAL1-sUbp1,* and *GAL1-sUbp1:RFP-NLS* constructs behaved as a *ubp111.* mutant. Total protein was extracted, resolved by 10% SDS-PAGE gels, and immunoblotted with anti-PCNA, anti-RFP, and anti-HA antibodies. PGK immunoblot was used as a loading control.

As mentioned above, Ubp1 has two different isoforms (35). We aimed to discriminate between these two forms in terms of their potential ability to regulate PCNA ubiquitylation. To this end, we replaced the endogenous *UBP1* promoter with the conditional *GAL1-10* promoter and engineered cells to express only one of the two isoforms by removing or not the 66 N-terminal amino acids (*GAL1-mUbp1* or *GAL1-sUbp1* strains, Figure 2A). In addition, these two Ubp1 forms were labeled with the fluorescent RFP epitope linked to a nuclear localization signal (NLS) to visualize their cellular localization. As a control, Ubp1 was also labeled with the RFP epitope linked to the NLS motif (Figure 2A). By fluorescence microscopy analysis we confirmed that while the larger Ubp1 variant (*GAL1-mUbp1-RFP:NLS)* surrounds the nucleus, probably co-localizing with the nuclear membrane, soluble Ubp1 (*GAL1-sUbp1-RFP:NLS* strain) shows a clear nuclear localization (Figure 2B). We then analyzed the effect of overexpression of these isoforms on the characteristic accumulation of ubiquitylated PCNA forms of *ubp1011. ubp1211.* mutant cells. As shown in Figure 2D, only an overexpression of the soluble isoform of Ubp1 counteracted the accumulation of Ub-PCNA (Figure 2D, compare lanes 7 and 8 in glucose versus galactose). Interestingly, we also found that the mere retention of endogenous Ubp1 in the nucleus (*Ubp1-RFP:NLS* strain), without protein overproduction, was able to restore wild-type levels of PCNA ubiquitylation in *ubp1011. ubp1211.* mutant cells (Figure 2D, lane 5). The same was observed in cells treated with MMS (0.02%) for 90 min to induce DNA damage-mediated ubiquitylation of PCNA. Retention of Ubp1 in the nucleus abolished the higher accumulation of ubiquitylated PCNA forms caused by the lack of both Ubp10 and Ubp12 PCNA-DUBs, reaching the levels of wild-type cells (Figure S3A). Moreover, the proliferation defects of *ubp1011. ubp1211.* mutant cells were also rescued in *ubp1011. ubp1211. ubp1:NLS* mutant cells (Figure S3B).

PCNA ubiquitylation dynamics in *ubp1011. ubp1211. ubp1-RFP:NLS* cells during the S phase were also analyzed (Figure 3A). As shown in Figure 3B, Ubp1-mediated PCNA deubiquitylation was effective and, consequently, no ubiquitylated forms of PCNA were detected during S phase progression, as occurs in wild-type cells. Interestingly, the delay in S phase progression observed in *ubp1011. ubp1211.* cells was abolished by Ubp1-mediated activity, which recovered a wild-type replication rate (Figure 3C). Moreover, the growth defects caused by the lack of Ubp10 and Ubp12 PCNA-DUBs were also rescued by the retention of Ubp1 in the nucleus (Figure 3D). These findings support the hypothesis that a soluble nuclear population of Ubp1 is involved in the regulation of PCNA deubiquitylation during DNA replication.

**Figure 3.**
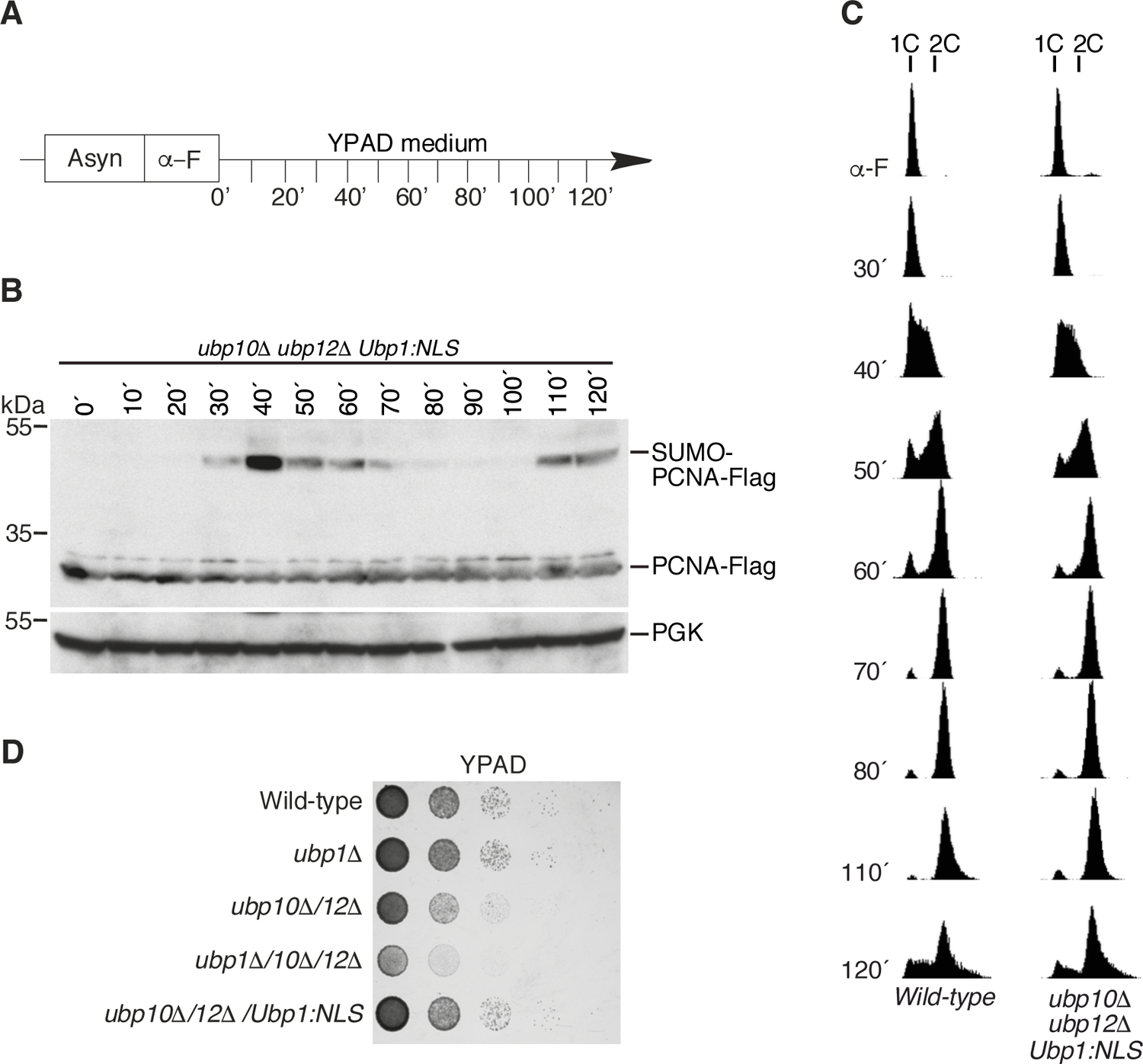
Retention of Ubp1 in the nucleus suppresses the S phase progression delay caused by the lack of the PCNA ubiquitin proteases Ubp10 and Ubp12 (A) Experimental design. Exponentially growing cultures of wild-type and *ubp1011. ubp1211. ubp1:NLS* strains expressing PCNA-Flag fusion protein were synchronized with α-factor and released into fresh medium (YPAD). Samples were taken at indicated intervals and processed for FACS and immunoblot analysis. (B) Protein extracts from *ubp1011. ubp1211. ubp1:NLS* strain were processed for immunoblotting with anti-Flag antibody. PGK protein was used as a loading control. (C) DNA content analysis by FACS shows the progression of genome replication from α-factor synchronization to the 120 minutes time point after release. Two biological replicates were performed, and a representative experiment is shown. (D) Ten-fold dilution assays of the indicated strains were incubated at 25 °C in a YPAD complex medium for 72 h and photographed.

### Ubp1 is a PCNA-DUB that associates with replication forks

The above observations correlated Ubp1 with PCNA deubiquitylation *in vivo*. Both the deletion and overexpression phenotypes of Ubp1 were consistent with the hypothesis that Ubp1 has a role as a PCNA ubiquitin protease. To further address this issue, we first checked whether nuclear Ubp1 is present in chromatin during DNA replication. Cells expressing Myc-tagged Ubp1 were synchronized with α-factor and then released into fresh YPAD medium to allow cells to progress through the S phase. Samples were taken at different time points and processed in order to obtain specific cellular fractions. Genome replication was followed by FACS. As shown in Figure 4A, although the vast majority of Ubp1 was detected in whole-cell extracts and chromatin-free fractions, it also appeared also linked to chromatin at all time points analyzed. These results correlate with those obtained by fluorescence microscopy (Figure 2B) and indicate that a nuclear population of Ubp1 is bound to chromatin.

**Figure 4.**
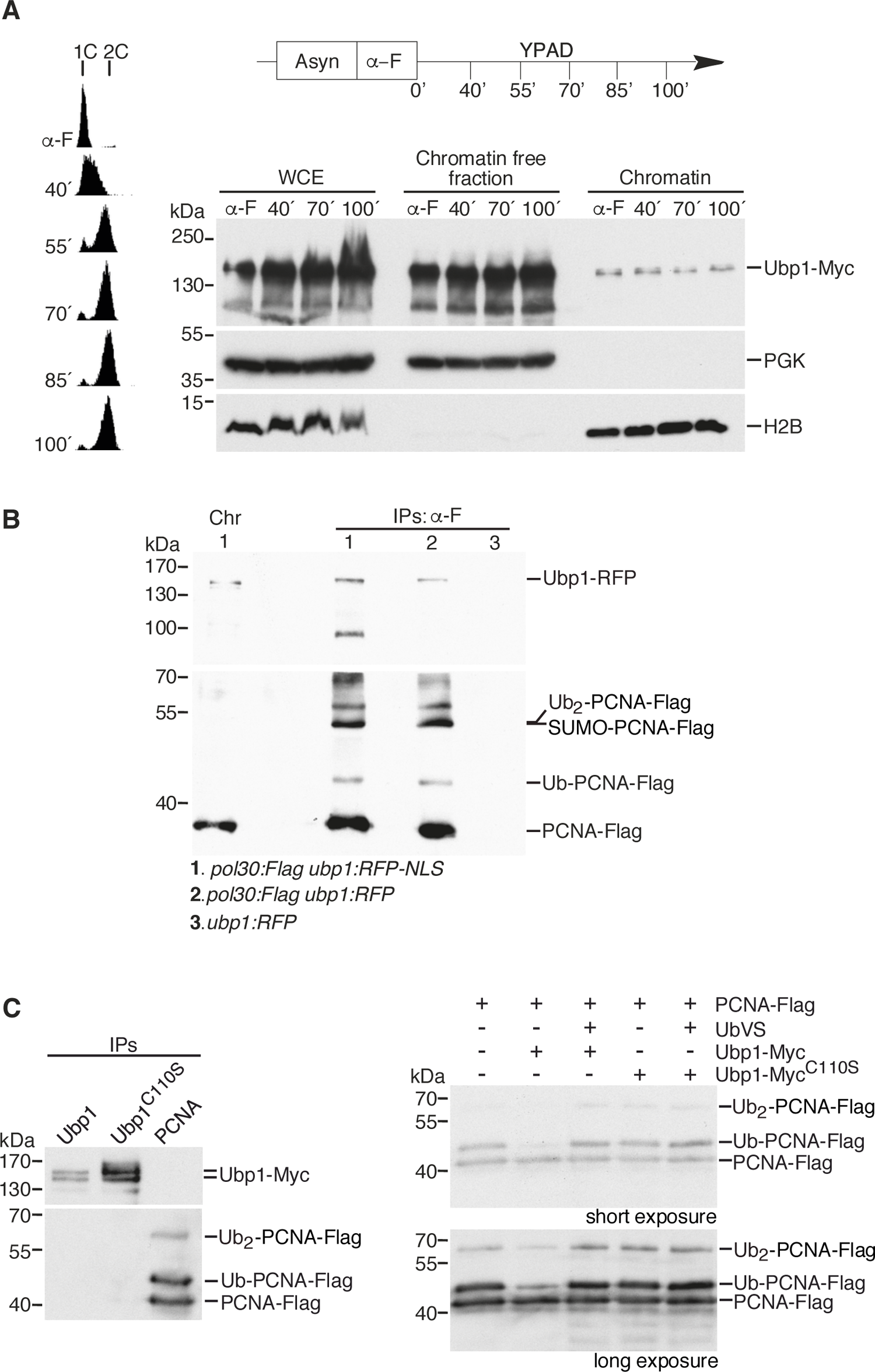
Ubp1 interacts *in vivo* with and deubiquitylates PCNA^K164^ (A) Ubp1 is associated with chromatin. Wild-type cells expressing Ubp1-Myc tagged protein were synchronized with α-factor and released into fresh yeast complex medium (YPAD) to allow progression through the S phase. Samples were taken at indicated time points and processed for FACS and immunoblot analyses. Soluble and chromatin protein fractions were resolved by SDS-PAGE gels, transferred to nitrocellulose membranes, and incubated with anti-Myc antibody. Histone H2B was used as a chromatin marker. WCE: whole cell extract. (B) Co-immunoprecipitation assay showing the interaction between PCNA-Flag and Ubp1-RFP fusion proteins *in vivo*. Cultures from the indicated strains were synchronized with α-factor and released in the presence of HU (0.2 mM) for 1 h. Samples were processed for ChIP and anti-Flag antibody were used to immunoprecipitate PCNA-Flag. Samples were resolved by SDS-PAGE gels and analyzed by immunoblotting. Cut blots were incubated with anti-Flag or anti-RFP antibodies. Chromatin extracts from *pol30:Flag Ubp1:RFP-NLS* were used as a reference for Ubp1-RFP detection. *ubp1:RFP* strain, no expressing PCNA-Flag was used as a negative control. Chr: chromatin, IP: immunoprecipitates. (C) PCNA *in vitro* deubiquitylation assay. Ubiquitylated PCNA was obtained by immunoprecipitation with anti-Flag antibody from an *S. pombe* strain expressing PCNA-Flag fusion protein and lacking *ubp12^+^*, *ubp15^+^* and *ubp16^+^*genes (left panel). Ubp1-myc or Ubp1^C110S^-Myc were also obtained by immunoprecipitation (left panel) from cells expressing each of these two proteins. PCNA immunoprecipitates were incubated with Ubp1-myc or Ubp1^C110S^-Myc in the absence or in the presence of Ub-VS, a ubiquitin protease inhibitor, for 1h at 30 °C. Samples were resolved by SDS-PAGE gels and analyzed by immunoblotting with anti-Flag antibody.

We next studied whether Ubp1 and PCNA interact *in vivo*. Since C-terminally RFP-tagged Ubp1 fusion proteins were found to be functional by fluorescence microscopy, we used these constructs in combination with Flag-PCNA recombinant protein. Thus, cultures of *pol30:Flag ubp1:RFP* and *pol30:Flag ubp1:RFP-NLS* strains were synchronized with α-factor and released for 1 hour in 0.2 M HU to slow S phase progression. Chromatin extracts were obtained and PCNA was immunoprecipitated with anti-Flag antibodies. We found that Ubp1 binds PCNA and that this binding increased when Ubp1 was retained in the nucleus (Figure 4B).

We then performed *in vitro* deubiquitylation assays to assess whether Ubp1 was able to directly deubiquitylate PCNA. Mono- and di-ubiquitylated PCNA was obtained by immunoprecipitation with anti-Flag antibodies from an *S. pombe* strain lacking the *ubp12^+^*, *ubp15^+^*, and *ubp16^+^* genes, which encode three of the four known PCNA-ubiquitin proteases in this organism, and also expressing PCNA-Flag fusion protein ((32) and Methods section). Ubp1-myc and Ubp1^C110S^-Myc, a catalytically inactive form of Ubp1, were immunoprecipitated from *S. cerevisiae* strains expressing each of these fusion proteins. As shown in Figure 4C, we found that Ubp1-myc exhibited remarkable activity on both Ub- and Ub_2_-PCNA forms. This activity was dependent on its catalytic residue cytosine 110 and was blocked by the irreversible DUB inhibitor ubiquitin vinyl sulfone (Ub-VS) (Figure 4C). All these data indicate that Ubp1 is a newly identified PCNA-DUB capable for removing ubiquitin moieties from both Ub- and Ub_2_-PCNA forms.

Ubp10 and Ubp12 PCNA-DUBs associate with replication forks while carrying out their function on PCNA (27). To understand if this is also the case for Ubp1 or, on the contrary, if Ubp1 can work apart from ongoing replication forks, the potential association of Ubp1 with different early replication origins was analyzed by ChIP-qPCR. Cells were synchronized by treatment with α-factor and released in the presence of HU (0.2 mM) for 1 hour to stall replication forks, and processed for Ubp1 ChIP, followed by qPCR analysis. Purified DNA samples were subjected to qPCRs using primers close to specific activated autonomous replicating sequences (ARSs) (e.g., *ARS305*, *ARS306*, *ARS603*, and *ARS607*). We found that Ubp1 associates with all the active replication origins analyzed (Figure 4D). We reasoned that the lack of Ubp10 and Ubp12 could favor this association. The binding of Ubp1 to replication origins in *ubp1011. ubp1211.* mutant cells was also analyzed. As shown in Figure 5, the recruitment of Ubp1 to all replication origins examined was independent of the presence or absence of Ubp10 and Ubp12 proteins. Altogether, these data indicate that Ubp1 participates in the regulation of PCNA ubiquitylation at replication forks independently of Ubp10 and Ubp12.

**Figure 5.**
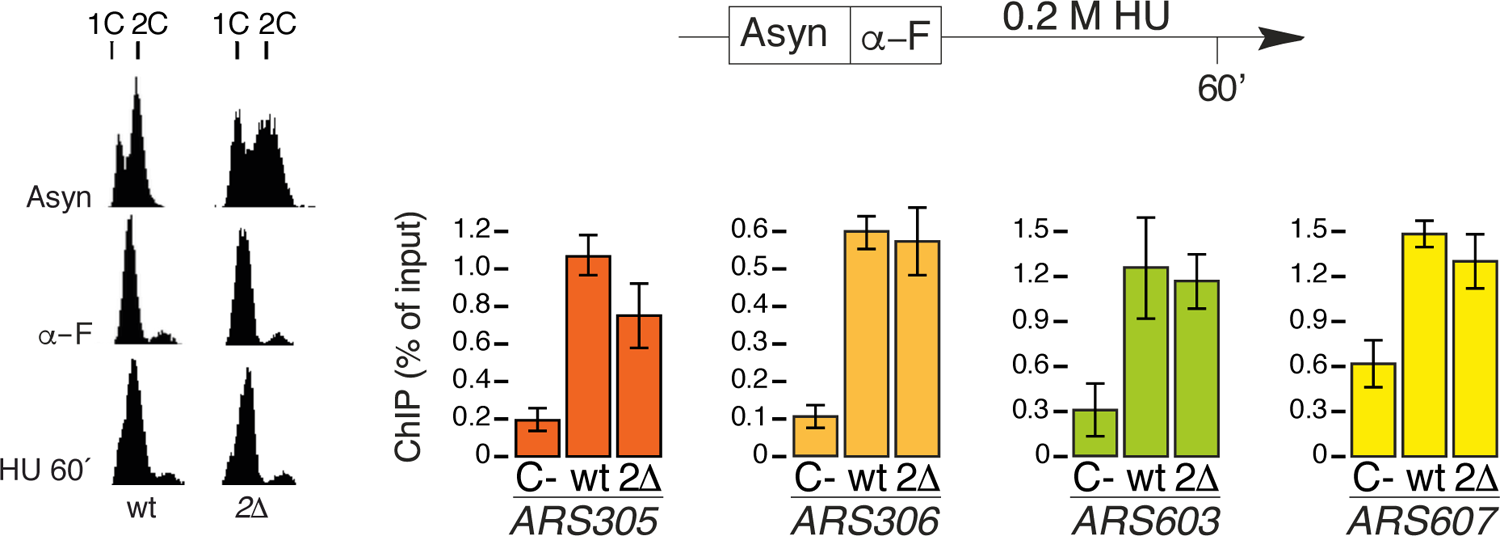
Ubp1 associates with replication forks Both wild-type and *ubp1011. ubp1211.* (*211.*) strains expressing Ubp1-Myc or wild-type untagged cells (C-) were synchronized with α-factor and released in the presence of HU (0.2 M). After 1 h, samples were collected and processed for ChIP-qPCR assays. Synchronization was confirmed by FACS. DNA content analysis corresponding to a particular experiment is shown on the left. qPCR reactions were performed by using specific primers for *ARS305*-, *ARS306*-, *ARS603*- and *ARS607*-proximal DNA fragment amplifications. Means and standard deviations for three independent experiments are shown.

### Ubp1 cooperates with Ubp10 and Ubp12 in the regulation of DDT processes

Ubp10 and Ubp12 PCNA-DUBs play a key role in the modulation of DDT during DNA replication by counteracting the engagement of nascent DNA strands in template switching (TS) events upon replication fork stalling. Therefore, the lack of these two DUBs exacerbates the accumulation of small Y-shaped intermediates to the detriment of the large Y-shaped ones, observed upon dNTP shortage induced by HU treatment in a rad52-dependent manner (27). We reasoned that if Ubp1 collaborated in PCNA deubiquitylation during S phase, it could also be involved in modulating the DDT response. To answer this question, we examined through neutral/neutral 2D gel electrophoresis (2D gels) the impact of *UBP1* depletion in combination with the double Ubp10/Ubp12 ablation on the replication intermediates pattern generated upon HU-induced fork stalling. As shown in Figure 6A-D, the lack of Ubp1 further increased the ratio of small/large Y-shaped intermediates observed in cells lacking Ubp10 and Ubp12 enzymes (27). It has been suggested that these small non-canonical Y-shaped replication structures most likely correspond to transitional structures in which the nascent DNA strands rearrange under these fork-stalling conditions. Moreover, when analyzing whether these structures corresponded to TS events, it was found that, as expected, they did. Thus, the accumulation of small Y-shaped molecules in the *ubp111. ubp1011. ubp1211.* mutant was suppressed by the depletion of Rad52 (Figure 6B-D). The increased accumulation of these Rad52-dependent replication intermediates in the *ubp111. ubp1011. ubp1211.* mutant compared to the double mutant *ubp1011. ubp1211.* following HU-induced replication fork stalling indicates that Ubp1 participates, along with Ubp10 and Ubp12 in the PCNA-DUB-driven TS branch of the DDT response at replication forks.

**Figure 6.**
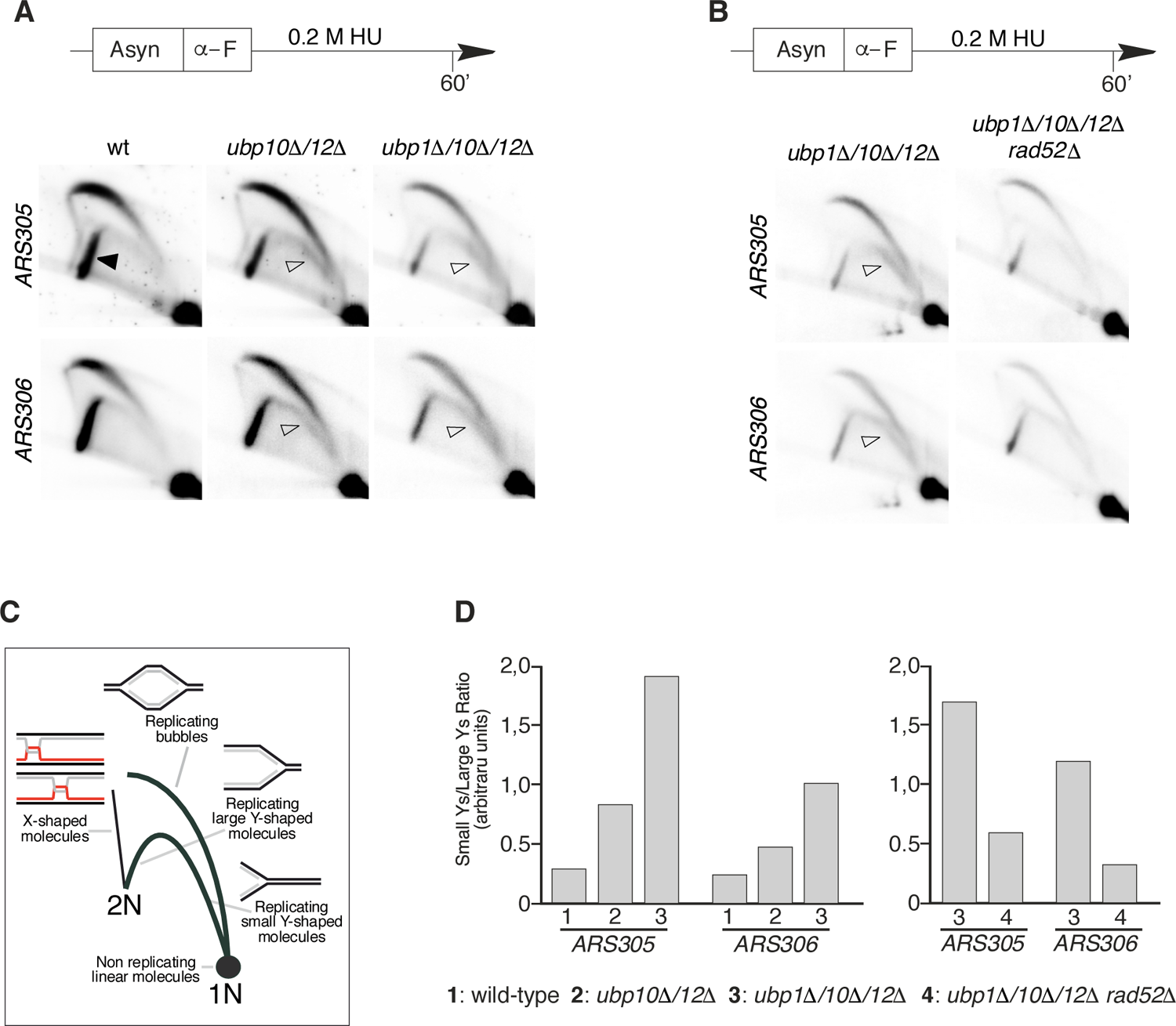
Accumulation of small Y-shaped TS intermediates at stalled forks in *ubp111. ubp1011. ubp1211.* mutant. (A, B) Wild-type, *ubp1011. ubp1211.* (*ubp1011./1211.*) and *ubp111. ubp1011. ubp1211.* (*ubp111. /1011. /1211.*) cells (A) or *ubp111. ubp1011. ubp1211.* and *ubp111. ubp1011. ubp1211. rad5211.* strains (B) were synchronized with α-factor and released in the presence of 0.2 M HU for 60 minutes. Samples were processed for FACS analysis of DNA content (Figure S4) and 2D gel analysis of replication intermediates. Genomic DNA was digested with NcoI endonuclease, resolved by 2D electrophoresis, transferred to nylon membranes, and hybridized to probes spanning *ARS305* and *ARS306* early replication origins. (C) Scheme of canonical replication intermediates and fully replicated joint molecules detected by 2D gel analysis. (D) Small/Large Y-shaped intermediate ratios are shown by histogram plots. Open and closed arrows indicate small and large Y-shaped structures, respectively, detected by 2D gel analysis.

## DISCUSSION

In this study, we provide a deep characterization of the *S. cerevisiae* ubiquitin protease Ubp1 in the context of DNA replication. Ubp1, hitherto only known as a cytoplasmic DUB, has a role as a nuclear protein in the deubiquitylation of PCNA at replication forks. In previous work, we revealed the functional impact of PCNA deubiquitylation in the control of DDT, and identified two specific ubiquitin proteases, Ubp10 and Ubp12, involved in it (27). In this research, by identifying a new PCNA-DUB, we confirm that a few deubiquitylating enzymes revert PCNA ubiquitylation during S phase in *S. cerevisiae* (27), as previously observed in *S. pombe* (32), suggesting that PCNA deubiquitylation is an important cellular process, in which cells use redundant enzymes to ensure the deubiquitylation of the PCNA sliding clamp. Underlining the importance of the removal of ubiquitin moieties from PCNA, the depletion of PCNA-DUBs generates a strong delay in S phase progression in both fission and budding yeasts ((27, 32) and this work), which is indicative of a role in the maintenance of processive DNA synthesis. One of the main activities of PCNA is to promote tolerance to DNA damage during DNA replication to prevent the formation of single-stranded DNA gaps, a potential cause of genomic instability and tumorigenesis (7, 14–16, 37, 38) and (39) for revision). Spontaneous DNA lesions, likely caused by endogenous cellular metabolic damage, may account for those dynamic PCNA ubiquitylation-mediated DDT mechanisms that prevent chromosome replication defects.

This study shows that the S phase progression defect observed in cells lacking Ubp10 and Ubp12 PCNA-DUBs (27) correlates with a transient accumulation of ubiquitylated PCNA during unperturbed DNA replication, not detected in wild-type cells. The transient nature of this pattern of PCNA ubiquitylation in the *ubp1011. ubp1211.* double mutant, which disappears at the end of the S phase, suggested that other ubiquitin protease(s) were also involved in the process. By generating triple mutants in which the lack of Ubp10 and Ubp12 was added to the loss of one of the 15 remaining specific ubiquitin proteases of the USP family known in *S. cerevisiae,* a third enzyme, Ubp1, participating in PCNA deubiquitylation was found. Consequently, ablation of Ubp1 in combination with Ubp10 and Ubp12 results in a much more pronounced delay in S phase progression and a permanent accumulation of ubiquitylated PCNA throughout DNA replication and beyond. Hence, the PCNA ubiquitinated levels conferred by MMS-mediated DNA damage were also higher in the *ubp111. ubp1011. ubp1211.* triple mutant compared to cells lacking only Ubp10 and Ubp12 proteins. In contrast, overexpression of Ubp1 in the nucleus provides the full rescue of both S phase progression and PCNA ubiquitylation defects observed in the triple mutant, strongly suggesting that Ubp1 is a PCNA-DUB.

The fact that only the ablation of Ubp10, but not of Ubp12 and Ubp1 by themselves, causes accumulation of ubiquitylated PCNA (33) suggests that all three enzymes collaborate in the deubiquitylation of PCNA; Ubp10 would play a major role, while Ubp12 and Ubp1 would contribute to a lesser extent, and Ubp12 and Ubp1 would take over some of the functions of Ubp10 when it is absent. Moreover, based on the pattern of PCNA ubiquitylation observed in *ubp111. ubp1011. ubp1211.* cells during unperturbed DNA replication, we could also hypothesize that Ubp1 could contribute to PCNA deubiquitylation by acting preferably at the end of S phase. It has been observed that Ubp10 and Ubp12 act differentially on PCNA, as Ubp10 efficiently removes all ubiquitin residues linked to PCNA^K164^, single ubiquitin monomers, or K^63^-linked polyubiquitin chains, whereas Ubp12 preferentially removes K63-linked ubiquitin moieties from poly-ubiquitylated molecules (27). Ubp1 was found to behave similarly to Ubp10, removing both single ubiquitin residues and poly-ubiquitin chains from PCNA. These different preferences for PCNA ubiquitin-chain removal suggest that Ubp10, Ubp12, and Ubp1 might cooperate in PCNA regulation by playing different roles in DNA damage bypass. Our findings reveal new levels of complexity in the PCNA-deubiquitylation-mediated DDT mechanisms, which still need further study.

Unexpectedly, we found that Ubp1 is a ubiquitin protease with a broader presence in cellular compartments beyond those known so far, the endoplasmic reticulum and cytoplasm (35). A subpopulation of Ubp1 located in the nucleus, where it associates with chromatin, was observed by fluorescence microscopy and biochemical analyses. Nuclear Ubp1 constitutes a small proportion of the total protein expressed in the cell, in particular of the soluble Ubp1 population. In addition, we show that nuclear Ubp1 interacts *in vivo* with PCNA and can remove both Ub- and Ub_2_-PCNA forms *in vitro*. This activity is also observed *in vivo* when driving Ubp1 to be fully expressed in the nucleus by adding an NLS to its C-terminal domain. The association of Ubp1 with replication forks is of particular interest, suggesting that, as in the case of Ubp10 and Ubp12 (27), the function of Ubp1 as a PCNA-DUB is carried out, at least in part, during replication fork progression.

Ubiquitylated PCNA accumulate when forks need to solve damaged templates or replicative stress conditions through DDT mechanisms (2, 7, 12, 13, 23). The fact that this accumulation is much higher in PCNA-DUB-deficient cells ((27); Figure 2C) indicates that deubiquitylation events are also involved. It has been shown that PCNA ubiquitin proteases downregulate DDT events acting at replication forks (27). Transient Rad52-dependent replication intermediates, which are quickly resolved in wild-type cells, accumulate in PCNA-DUB-deficient cells (27). During replication of alkylated DNA, X-shaped TS intermediates accumulate due to nascent strand exchange events, leading to the formation of joint molecules that eventually dissolve in wild-type cells (25, 40, 41). In Ubp10- and Ubp12-deficient cells these Rad52-dependent molecules are not efficiently resolved and accumulate over longer periods, probably due to increased PCNA ubiquitylation (27). Similarly, under replicative stress conditions generated by dNTPs shortage, the accumulation of small non-canonical Y-shaped Rad52-dependent structures was also observed in Ubp10- and Ubp12-deficient cells but not in the wild-type cells. These anomalous replicative intermediates accumulate at the expense of the large canonical Y-shaped ones, probably due to an incomplete synthesis of the nascent strands (27). Here we show that in cells lacking Ubp1 in combination with Ubp10 and Ubp12 ablation, the ratio of small/large Y-shaped intermediates generated after HU treatment is higher than that observed in *ubp1011. ubp1211.* mutant cells and that this phenotype depends on Rad52, which strongly suggests that Ubp1 is also involved in TS-mediated DDT modulation mechanisms at replication forks.

Since the presence of Ubp1 at replication forks is not affected by the deletion of Ubp10 and Ubp12, we can hypothesize that there is not regulation between them and that they could act on PCNA independently. How these three enzymes work together in these DDT processes, what the function of each enzyme is, where and when each enzyme carries out its work, and how they are regulated are very interesting open questions to address in future studies.

Many factors involved in replication and the response to replicative stress are known to be regulated by ubiquitylation (6, 42–44). Since Ubp1 associates with chromatin not only during early S phase but throughout the interphase, we cannot rule out the possibility that its contribution to DNA replication could be also carried out through additional substrates other than PCNA. Ubp7, initially characterized as an endocytic factor in *S. cerevisiae*, has also been proposed as a component of the regulatory network of S phase progression under conditions of DNA damage, probably working on the chromatin state through a so-far unknown substrate (45). Additionally, Ubp1 could be involved in additional nuclear functions other than DDT and DNA replication.

These results indicate that deubiquitylation-dependent regulation of DNA replication is a complex network involving several ubiquitin-proteases. How the different PCNA-DUBs work and are regulated remains unknown. Molecular mechanistic studies will shed light on PCNA deubiquitylation-dependent processes. Genome instability constitutes a hallmark of age-related diseases such as cancer (46, 47), and it is during DNA replication that cells are most vulnerable to losing genomic stability. A better understanding of the molecular mechanisms that regulate DNA replication is the basis for both a better knowledge of cancer and the development of more precise therapies.

## MATERIAL AND METHODS

### Yeast strains, growth conditions and media

All the budding yeast used in these studies originate from a *MATa* W303 *RAD5 bar1::LEU2* strain (33) and are listed in the Supplementary information Table S1. For the *in vitro* analysis of Ubp1 activity, a fission yeast strain listed in the Supplementary information Table S1 was used as a source of ubiquitylated PCNA. Budding yeast strains were grown in YPAD medium (1% yeast extract, 2% peptone supplemented with 50 μg/ml adenine) containing 2% glucose. For block-and-release experiments, cells were grown in YPAD with 2% glucose at 25°C and synchronized in G1 with α-factor pheromone (40 ng/ml, 2.5 hours). Cells were then collected by centrifugation (3000 rpm 3 min) and released into fresh media (supplemented with 50 µg/ml of pronase) in the absence or in the presence of HU (0.2 M, FORMEDIUM). Overexpression experiments with cells grown in YPAD medium with 2% raffinose at 25°C were conducted by adding to the medium 2.5% galactose (to induce) or 2% glucose (to repress).

For plate survival assays stationary cells were counted and serially diluted in YPAD media. Ten-fold dilutions of equal numbers of cells were plated onto YPAD (2% glucose) media (always supplemented with 50 μg/ml adenine), or YPAD containing 0.02% MMS, incubated at 25°C for 24, 48, 72 or 120 hours and then scanned.

### General experimental procedures

General experimental procedures of yeast Molecular and Cellular Biology were used as described previously (48–50). A list of the plasmids used for strains generation is shown in the Supplementary Table S2. Transformation was performed by lithium acetate protocol and transformants were selected by growing in selective medium.

### Flow Cytometry Analysis

For flow cytometry analyses, 10^7^ cells were collected by centrifugation, washed once with water, fixed in 70% ethanol and processed as described previously (51). Cells were prepared using a modification of the method, by using SYTOX Green (Molecular PROBES) for DNA staining (52, 53). The DNA content of individual cells was measured using a Becton Dickinson Accuri C6 plus FACScan.

### Protein Methods

#### Protein Extracts Preparation and Western blot analysis

Whole cell extracts were prepared by precipitation with trichloroacetic acid (TCA). Yeast strains were grown in YPAD medium to OD_600_ of 0.8-1.0 and cells (5 ml) were collected by centrifugation just after the addition of 100% TCA to a final concentration of 10% TCA and washed with 20% TCA. Cell disruption was performed with Glass-Beads in a Fast-Prep and 12.5% TCA. Cell lysates were pelleted by centrifugation at 3000 rpms and resuspended in 1X LB loading buffer and Tris base.

For chromatin-enriched fractions around 6 x 10^7^ exponentially growing cells were harvested by centrifugation and resuspended in 1 ml of Buffer 1 (containing 150 mM Tris pH 8.8, 10 mM dithiothreitol (DTT), and 0.1% sodium azide), and incubated at room temperature for 10 minutes. Cells were pelleted, washed with 1 ml of Buffer 2 (50 mM KH_2_PO_4_/K_2_HPO_4_ pH 7.4, 0.6 M Sorbitol, and 10 mM DTT), resuspended in 200μl of Buffer 2 supplemented with 40 μg Zymolyase-100T and incubated at 37 °C for 10 minutes with intermittent mixing. The resulting spheroplasts were washed with 1 ml of ice-cold Buffer 3 (50 mM HEPES pH 7.5, 100 mM KCl, 2.5 mM MgCl, and 0.4 M Sorbitol), followed by resuspension and a 5-minute incubation in 100 μl of EBX buffer (50 mM HEPES pH 7.5, 100 mM KCl, 2.5 mM MgCl, 0.25% Triton100, 1 mM phenylmethylsulfonyl fluoride (PMSF), Protease inhibitor tablets (EDTA-free, Roche), Leupeptin 1 μg/ml, Pepstatin 2.5 μg/ml, and RNAse 10 μg/ml), with occasional mixing. Aliquots of 30 μl of these disrupted cell suspensions were collected as whole cell extract samples (WCE). Remaining volume was layered onto 70 μl of cold EBX-S buffer (EBX buffer supplemented with 30% Sucrose) and subjected to centrifugation at 12000 rpm for 10 minutes at 4 °C. Aliquots of 30 μl of the resulting supernatant layer (Chromatin-free fraction) were also collected. After discarding supernatant, chromatin pellets were washed with 200 μl of EBX-S buffer, resuspended in 70μl of EBX buffer supplemented with 0.5 μl of Benzonase, and incubated on ice for 15 minutes (Chromatin fraction). 5X loading buffer was added to each fraction.

Protein extracts were electrophoretically resolved by SDS-PAGE (8%, 10%, 12% or 15%) gels and transferred to Nitrocellulose membranes using a Bio-Rad transfer unit. Blots were then probed against antibodies indicated. A list of the antibodies used in this study is shown in the Supplementary Table S3. Secondary horseradish peroxidase-conjugated anti-rabbit, anti-goat, or anti-mouse antibodies (as required) were also used and the ECL kit (Amersham Pharmacia Biotech) for detection.

#### Co-Immunoprecipitation

Immunoprecipitation of Flag-tagged PCNA protein was performed from chromatin extracts of strains expressing PCNA-Flag and/or Ubp1-RFP fusion proteins. Cells were grown in YPAD medium at 25 °C to an OD_600_ of 0.8-1.0 (25 ml), synchronized with α-factor and released in the presence of HU (0.2 M) for 90 min. Chromatin extracts were prepared as indicated in the “*ChIP-qPCR analysis*” section of Material and Methods. Extracts were incubated with dynabeads protein G (Invitrogen) bound to monoclonal anti-Flag antibody (Agilent Technologies) for 5 hours at 4°C. Beads were washed four times with lysis buffer and resuspended in loading buffer. Immunoprecipitates were resolved by SDS-PAGE gels, transferred to Nitrocellulose membranes and analysed with anti-RFP (Chromotek) and anti-Flag-HRP conjugated (Sigma) antibodies.

#### *In vitro* deubiquitylation assays

Immunoprecipitation of PCNA-FLAG was performed from a *ubp12*-NES *ubp15*-NES *Δubp16 pcn1-FLAG S. pombe* strain (see Supplementary Table S1), synchronized in S phase by treatment with 20 mM HU for 2 hours. *S. pombe* PCNA is a reliable and abundant source of ubiquitylated PCNA lacking SUMO-PCNA, which would otherwise hamper our *in vitro* assay (32). Immunoprecipitation of Myc-tagged Ubp1 proteins were performed from asynchronously growing cells. Both wild-type and the catalytically inactive form of Ubp1, were purified from soluble protein extracts prepared as described previously (54). Briefly, cells were collected, washed, and broken in HB2T buffer (60 mM β-glycerophosphate, 15 mM *p*-nitrophenylphosphate, 25 mM 4-morpholinepropanesulfonic acid (pH 7.2), 15 mM MgCl2, 15 mM EGTA, 1 mM DTT, 0.1 mM sodium orthovanadate, 2% Triton X-100, 1 mM PMSF, and 20 mg/ml leupeptin and aprotinin) using glass beads. Glass beads were washed with 500 μL of HB2T, and supernatant was recovered. Protein concentrations were measured using the BCA assay kit (Pierce) and immunoprecipitations (from 4 mg of protein extracts) were carried out by incubation with anti-Myc or anti-Flag antibodies-bound magnetic beads during 5 hours at 4 °C. Immunoprecipitation assays were confirmed by immunoblotting. Immunoprecipitates were washed twice with lysis buffer and then twice with DUB buffer (60 mM HEPES pH 7.6, 5 mM MgCl2, 4% glycerol). Beads were incubated 1 hour at 30°C. As negative controls, catalytically inactive Ubp1 (Ubp1^C110S^) was used. Vinyl sulfone (Ub-VS) (Enzo Life Sciences) covalently captures active DUB enzymes and therefore acts as a potent and irreversible inhibitor of DUBs through the covalent modification of their active sites (55). Reactions were stopped by adding loading buffer and boiling the samples for 5 min at 95°C. Proteins were resolved by SDS-PAGE gels, transferred to nitrocellulose membranes and analyzed with anti-Flag-HRP conjugated antibody.

#### ChIP-qPCR analysis

We adapted a described protocol (43) for the analysis of myc-tagged Ubp1 *ARS305, ARS306, ARS603* or *ARS607* binding in *S. cerevisiae* cells. In brief, Ubp1-Myc or wild-type untagged cells (used as control) were synchronized with α-factor and released in the presence of HU (0.2 M). After 1 hour, samples (50 ml cultures) were taken and subjected to 30 minutes of crosslinking with 1% formaldehyde. Then, cells were collected by centrifugation and washed three times with ice-cold TBS. Cell pellets were resuspended in Lysis Buffer (50 mM Hepes pH 7.5, 140 mM NaCl, 1 mM EDTA, 1% Tritón-X100, 0.1% Na-deoxycholate) supplemented with Antiproteolytic Cocktail and broken using glass beads.

Recovered cell lysates were centrifugated at 12000 rpm, supernatants (soluble protein fractions) were discarded, and chromatin pellets were sheared by sonication. Extracts were clarified and soluble chromatin fractions were used for immunoprecipitation with anti-Myc antibodies (5 hours at 4°C). Antibody-bound magnetic beads were washed as for CoIPs assays and chromatin was eluted in Elution Buffer (50 mM Tris pH 8.0, 10 mM EDTA, 1% SDS) by incubating 10 min at 65°C. Samples were incubated overnight at 65°C in TE (+1% SDS) for de-crosslinking, treated with Proteinase K, DNA extracted by phenol/chlorophorm/isoamylalcohol pH 8.0 and treated with 0.3 μg/ml RNase A in TE. Finally, DNA was purified with QIAquick® PCR purification kit and 1-10 ng of immunoprecipitated or input DNA were amplified with iQ™ SYBR Green Supermix (BioRad) using a Real-Time PCR machine (BioRad IQ™ 5). A list of the specific primers used is shown in the Supplementary Table S4. All data in the bar graphs are presented as an average of n ≥ 3 replicates ± standard deviation (SD), where n represents the number of biological replicates.

#### Two-dimensional DNA gels (2D-gel analysis)

DNA samples for neutral-neutral two-dimensional gel electrophoresis were prepared and analyzed as described previously (48, 56). DNA was cut with the NcoI restriction enzyme, transferred to Hybond-XL (GE Healthcare) nitrocellulose membrane and hybridized to probes spanning the *ARS305* and *ARS306* origins of DNA replication. For each origin of replication tested, the specific probe corresponds to the following coordinates (retrieved from SGD): *ARS305* (39073-40557, Chr III) and *ARS306* (73001-73958, Chr III). Images were acquired using a Molecular Imager FX (BioRad) and different replication-associated DNA molecules were quantified using Quantity One 4.6 software (BioRad).

#### Mycroscopy

GFP- and RFP-tagged strains were grown in YPAD or YPA + 2,5% galactose medium until exponential phase. Expression of the different chimeric Ubp1 forms was either repressed by adding glucose or induced with galactose in the medium. DAPI staining was used to visualize DNA and the presence of specific fluorescence was detected by fluorescence microscopy using a Thunder Imager 3D Tissue (camera, DFC9000; Leyca) microscope.

## ACKNOWLEDGMENTS

We are grateful to members of 08 group at the IBMCC and R. Bermejo for helpful discussions. This work was supported by the Spanish Ministry of Science (Grant reference PID2019-109616GB-100 to A.B. and M.P.S.) and Junta de Castilla y León (Grant reference SA103P20 to A.B). J.Z. was supported by a predoctoral fellowship from the Junta de Castilla y León. A.B. and M.P.S. Institution is supported by the “Programa de Apoyo a Planes Estratégicos de Investigaciόn de Excelencia” cofunded by the Junta de Castilla y Leόn and the European Regional Development Fund (CLC-2017-01).

## Author Contributions

Conceptualization, A.B. and M.P.S.; Investigation, J.Z.; Supervision, A.B. and M.P.S.; Formal Analysis, A.B., M.P.S. and J.Z.; Writing & Editing, M.P.S. and A.B.; Funding Acquisition, A.B. and M.P.S.

