## Supplemental Information for "Ubiquitin protease Ubp1 cooperates with Ubp10 and Ubp12 to revert Lysine-164 PCNA ubiquitylation at replication forks"

**Supplementary Information**

**Supplementary Figure S1**

Analysis of ubiquitylated PCNA levels of the indicated strains growing under unperturbed conditions. Total protein extracts were extracted from asynchronously growing cells expressing Flag-tagged fusion protein, resolved by 10% SDS-PAGE gels and analyzed by immunoblotting with anti-Flag antibody. Ponceau immunostaining was used as a loading control.

**Supplementary Figure S2**

Analysis of PCNA by immunoblotting using a rabbit polyclonal antibody that specifically detects PCNA monoubiquitylated forms in yeast cell extracts. Immunoblot of protein extracts from wild-type, *rad18*Δ (unable to ubiquitylate Lysine 164 PCNA), *mms2*Δ (unable to poly-ubiquitylate PCNA), $pol30^{K164R}$ (unable to ubiquitylate or SUMOylate Lysine 164 PCNA), and the *ubp1*Δ *ubp10*Δ *ubp12*Δ triple mutant (which accumulate high levels of ubiquitylated $PCNA^{K164}$) cells treated 90 minutes with 0.02% MMS. Ponceau immunostaining was used as a loading control.

**Supplementary Figure S3**

(A) Immunoblot analysis of ubiquitylated PCNA levels of the indicated strains. Asynchronously growing cells (untreated) were treated with 0.02% MMS for 90 min. Total protein extracts were extracted, resolved by 10% SDS-PAGE gels and immunoblotted with anti-Flag antibody. Ponceau immunostaining was used as a loading control. (B) Ten-fold dilution assays of the indicated strains were spotted on YPAD solid plates containing or not 0.02% MMS, incubated at 25 ºC for the indicated times, and photographed. Representative experiments are shown.

**Supplementary Figure S4**

(A, B) Experimental design at top. Exponentially growing cultures of the indicated strains were synchronized at G1 by incubation with α-factor and then released into complex medium (YPAD) containing 0.2 M HU for 1 hour. Samples were taken at indicated time points and processed for FACS analysis.

**Supplementary Table S1**

| Genotype | Strain | Source |
|---|---|---|
| *Saccharomyces cerevisiae* W303 Mata ade2-1 can1-100 his3-11,15 leu2-3,112 trp1-1 ura3-1 RAD5$^{+}$ bar1::LEU2 | 55.34 | Lab stock |
| 55.34 with Pol30:3Flag:kanMX | 61.45 | Lab stock |
| 55.34 with Pol30:3Flag:kanMX ubp10::natMX | 62.05 | Lab stock |
| 55.34 with Pol30:3Flag:kanMX ubp10::natMX ubp12::hphMX | 71.54 | Lab stock |
| 55.34 with Pol30:3Flag:kanMX ubp10::natMX ubp12::hphMX ubp1::HIS3 | 81.69 | This study |
| 55.34 with Pol30:3Flag:HIS3 ubp10::natMX ubp12::hphMX Ubp1:RFP-NLS:kanMX | 84.08 | This study |
| 55.34 with Ubp1:GFP:kanMX | 82.35 | This study |
| 55.34 with Ubp1:GFP-NES:natMX ubp10::hphMX ubp12::kanMX | 82.51 | This study |
| 55.34 with HIS3:Gal1:10:3Ha:mUbp1 ubp10::hphMX ubp12::kanMX | 83.32 | This study |
| 55.34 with HIS3:Gal1:10:3Ha:mUbp1:RFP-NLS:kanMX ubp10::natMX ubp12::hphMX | 83.12 | This study |
| 55.34 with HIS3:Gal1:10:3Ha:sUbp1 ubp10::natMX ubp12::hphMX | 83.38 | This study |
| 55.34 with HIS3:Gal1:10:3Ha:sUbp1:RFP-NLS:kanMX ubp10::natMX ubp12::hphMX | 83.48 | This study |
| 55.34 with Ubp1:RFP-NLS:kanMX ubp10::natMX ubp12::hphMX | 83.23 | This study |
| 55.34 with ubp1::HIS3 | 82.49 | This study |
| 55.34 with ubp10::hphMX | 61.33 | Lab stock |
| 55.34 with ubp10::hphMX ubp12::kanMX | 69.05 | Lab stock |
| 55.34 with ubp10::HIS3 ubp12::kanMX ubp1::natMX | 75.72 | Lab stock |
| 55.34 with Pol30:3Flag:HIS3 Ubp1:RFP:kanMX | 87.40 | This study |
| 55.34 with Pol30:3Flag:HIS3 Ubp1:RFP-NLS:kanMX | 87.41 | This study |
| 55.34 with Ubp1:RFP:kanMX | 87.18 | This study |
| 55.34 with Ubp1:13myc:kanMX | 83.10 | This study |
| 55.34 with ubp10::natMX ubp12::hphMX Ubp1:13myc:kanMX | 83.44 | This study |
| 55.34 with ubp10::HIS3 ubp12::kanMX ubp1::natMX $Ubp1_{c110s}$:13myc:URA3 | 83.33 | This study |
| 55.34 with ubp10::HIS3 ubp12::kanMX ubp1::natMX rad52::URA3 | 89.01 | This study |
| *Schizosaccharomyces pombe* h-; pcn1:3FLAG:KanMX6; ubp12:GFP-NES:NatMX4; ubp15:mRFP-NES:KanMX6; ubp16::HphMX4; ade6-M210; leu1-32; ura4D18 | 73.69 | Lab stock |

**Supplementary Table S2**

| Plasmids |
|---|
| pFA6a:3Flag:HIS3 |
| pFA6a:RFP:kanMX |
| pFA6a:RFP-NLS:kanMX |
| pFA6a:GFP:kanMX |
| pFA6a:GFP-NES:natMX |
| pFA6a:HIS3:Gal1-10:3HA |
| pFA6a:13MYC:kanMX |
| pRK983: Ubp1:13myc C110S:URA3 (From Dr. Kölling R.) |
| pBluescript:KlURA3 |

**Supplementary Table S3**

| Antibodies | Source and identifier |
|---|---|
| Anti-Flag M2 peroxidase (HRP) | Sigma-Aldrich (*Ref: A8592-1MG)* |
| Anti-PCNA | Generated at home |
| Anti-Rad53 (yN-19) | Santa Cruz Biotechnologies (*Ref: Sc-6748)* |
| Anti-HA-HRP | Miltenyi Biotec (*Ref: 130-091-972)* |
| Anti-PGK | Molecular Probes (*Ref: A-6457)* |
| Anti-Histone H2B | Active Motif (*Ref: 38237)* |
| Anti RFP | Chromotek (*Ref: 5f8)* |
| Anti c-Myc (HRP) | Miltenyi Biotec (*Ref: 130-092-113)* |
| Anti c-Myc | Sigma Aldrich (Ref: M5546-2Ml) |
| Anti-rabbit IgG-Peroxidase | Amserham (Ref: NA934) |
| Anti-mouse IgG-Peroxidase | Amserham (Ref: NA931) |

**Supplementary Table S4**

| Primers | Sequences |
|---|---|
| ARS 305 Forward | GTA ACT TAC ACG GGG GCT AA |
| ARS 305 Reverse | ACT TTG ATG AGG TCT CTA GC |
| ARS 306 Forward | GGA CAA GGT GCA AAT GCC AAG |
| ARS 306 Reverse | CCC GCT CCT TCT CCT AAA CAT |
| ARS 603 Forward | CAA ACC ACC GTC AAC ACC |
| ARS 603 Reverse | CCT CGA GGG TCG AAA TC |
| ARS 607 Forward | CAC ATT ATT CGG CAC AGT AGG |
| ARS 607 Reverse | GTG TCG CAG TCC ATA GAA GG |

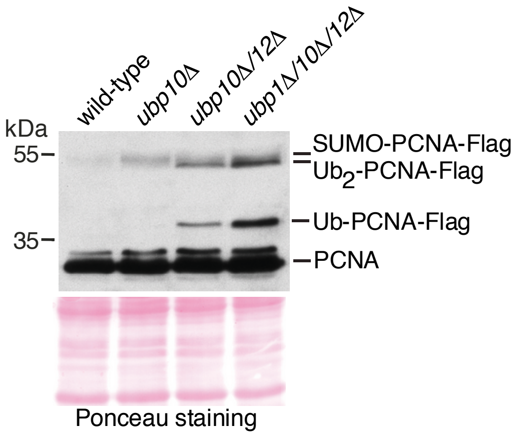
wild-type
ubp10∆
ubp10∆/12∆
ubp1∆/10∆/12∆
kDa
55
35
SUMO-PCNA-Flag
Ub2-PCNA-Flag
Ub-PCNA-Flag
PCNA
Ponceau staining

Zamarreño et al. Figure S2

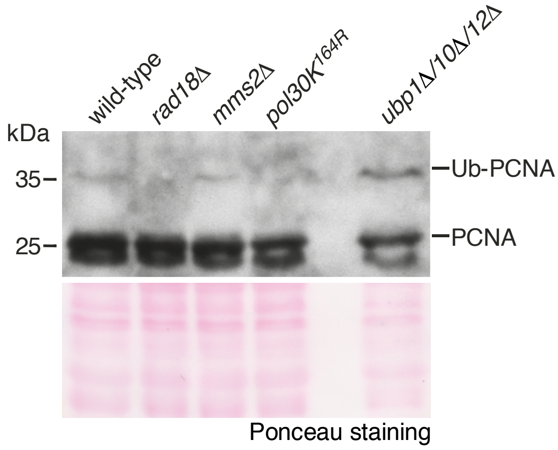

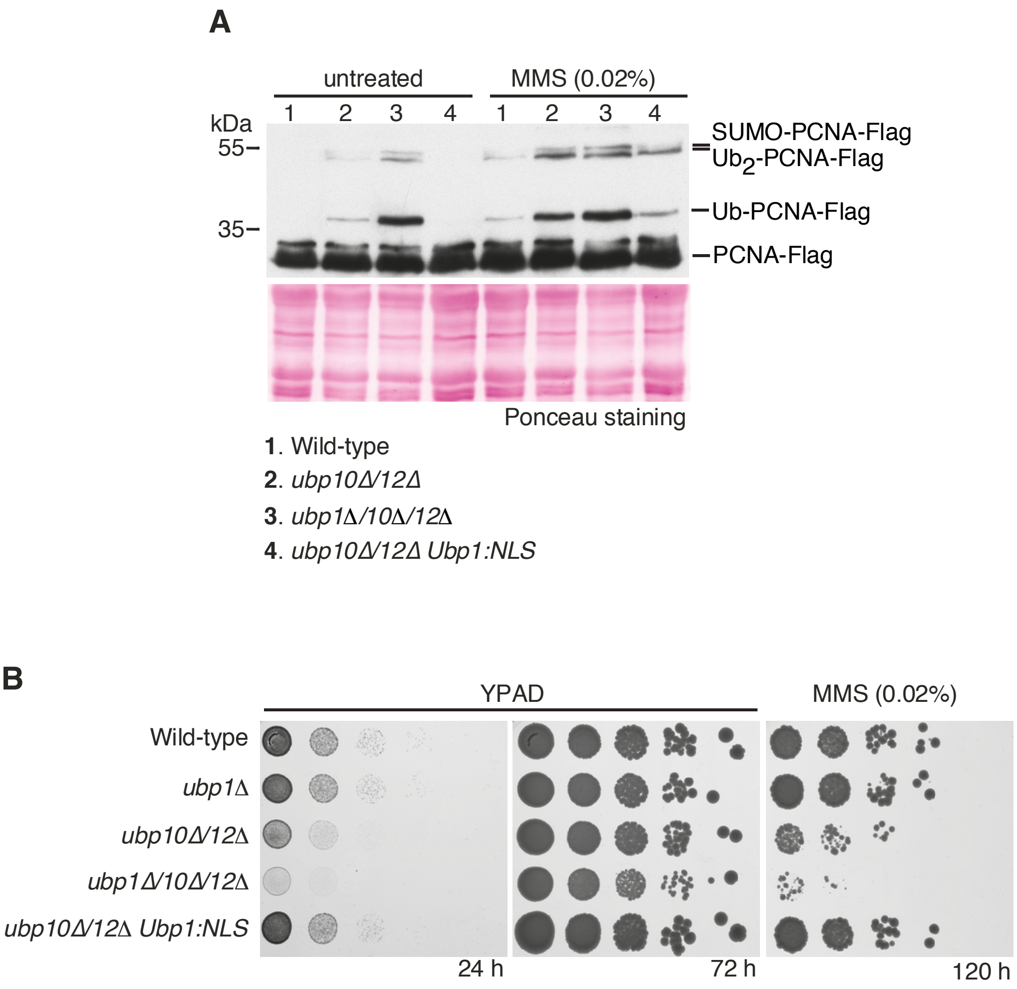

A
untreated
MMS (0.02%)
1 2 3 4 1 2 3 4
kDa
55–
35–
SUMO-PCNA-Flag
Ub2-PCNA-Flag
Ub-PCNA-Flag
PCNA-Flag
Ponceau staining
1. Wild-type
2. ubp10Δ/12Δ
3. ubp1Δ/10Δ/12Δ
4. ubp10Δ/12Δ Ubp1:NLS
B
YPAD
MMS (0.02%)
Wild-type
ubp1Δ
ubp10Δ/12Δ
ubp1Δ/10Δ/12Δ
ubp10Δ/12Δ Ubp1:NLS
24 h
72 h
120 h

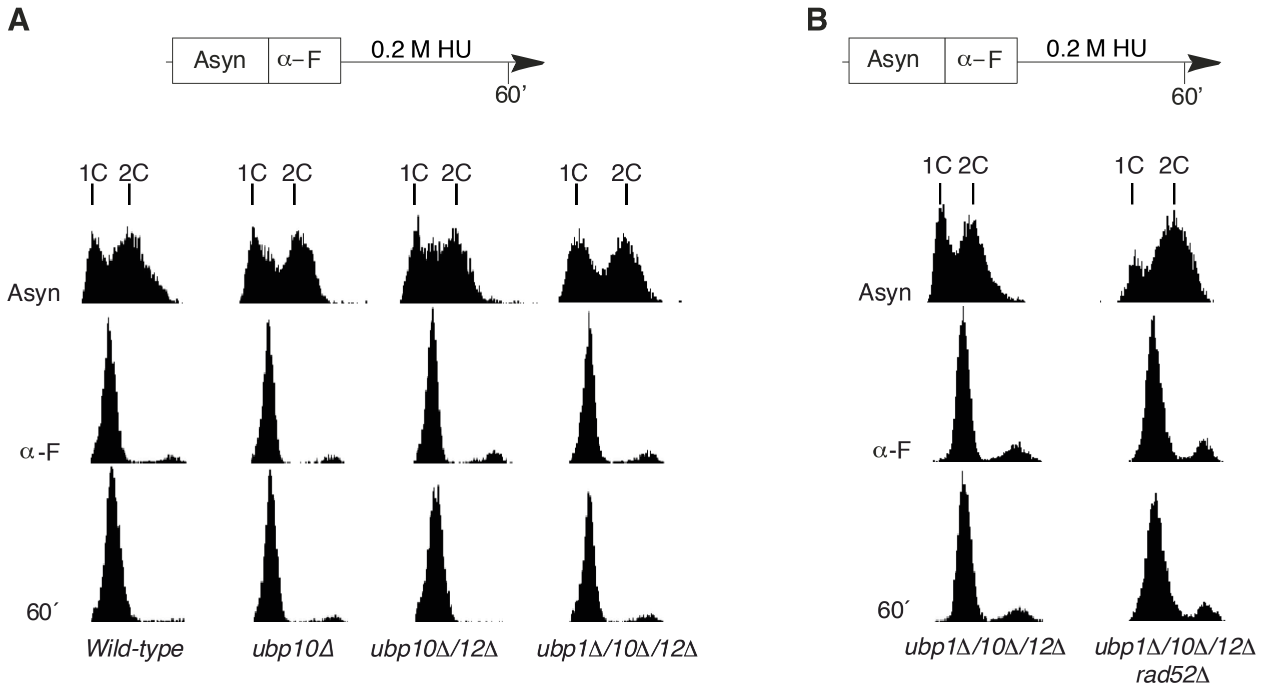
A
Asyn
α–F
0.2 M HU
60’
1C 2C
Asyn
α-F
60´
Wild-type
ubp10Δ
ubp10Δ/12Δ
ubp1Δ/10Δ/12Δ
B
Asyn
α–F
0.2 M HU
60’
1C 2C
Asyn
α-F
60´
ubp1Δ/10Δ/12Δ
ubp1Δ/10Δ/12Δ rad52Δ
